# How to limit the speed of a motor: The intricate regulation of the XPB ATPase and Translocase in TFIIH

**DOI:** 10.1101/2020.08.24.264176

**Authors:** J. Kappenberger, W. Koelmel, E. Schoenwetter, T. Scheuer, J. Woerner, J. Kuper, C. Kisker

## Abstract

The superfamily 2 helicase XPB is an integral part of the general transcription factor TFIIH and assumes essential catalytic functions in transcription initiation and nucleotide excision repair. In both processes the ATPase activity of XPB is essential. We investigated the interaction network that regulates XPB via the p52 and p8 subunits with functional mutagenesis based on a crystal structure of the full p52/p8 complex and current cryo-EM structures. Importantly, we show that XPB’s ATPase can be activated either by DNA or by the interaction with the p52/p8 proteins. Intriguingly, we observe that the ATPase activation by p52/p8 is significantly weaker than the activation by DNA and when both p52/p8 and DNA are present, p52/p8 dominates the maximum activation. We therefore define p52/p8 as the master regulator of XPB that acts as an activator and speed limiter at the same time. A correlative analysis of the ATPase and translocase activities of XPB shows that XPB only acts as a translocase within the context of complete core TFIIH and that XPA increases the processivity of the translocase complex without altering XPB’s ATPase activity. Our data unravel an intricate network that tightly controls the activity of XPB during transcription and nucleotide excision repair.

## Introduction

The ATPase activity of the superfamily 2 (SF2) helicase XPB is indispensable for two main cellular processes, transcription and nucleotide excision repair (NER) (1-4). XPB is part of the general transcription factor IIH (TFIIH) (5-7), which is functionally divided into core TFIIH and the CAK complex. Besides XPB, the core complex contains a second SF2 helicase, XPD, as well as the p62, p52, p44, p34 and p8 subunits. The CAK (cyclin-dependent-kinase (CDK)-activating kinase) complex completes the TFIIH architecture and consists of MAT1, Cyclin H and the kinase CDK7. Holo-TFIIH orchestrates the three differential enzymatic activities of XPB, XPD and CDK7. The individual roles of the three enzymes present in TFIIH differ depending on the cellular processes that TFIIH is involved in. During RNA polymerase II (RNAPII)-based transcription the ATPase activity of XPB is required for promotor opening, while XPD solely assumes a scaffolding function (6, 8-11). Phosphorylation of the C-terminal domain (CTD) of the Rpb1 subunit of RNAPII by the CAK subcomplex promotes the initiation and elongation in transcription (12-15), whereas the same complex assumes an inhibitory effect on the NER activity and was shown to dissociate from core TFIIH prior to the incision of the damaged strand (16, 17). In contrast to transcription initiation, both the ATPase activity of XPB and the ATP-dependent helicase activity of XPD are essential for the NER pathway to anchor TFIIH to the site of the damaged DNA and to open the DNA duplex in the context of the lesion (6, 8-11). The significance of an intact TFIIH complex for NER is reflected in its association with three severe autosomal recessive disorders. Mutations in the *ERCC3* (XPB) or *ERCC2* (XPD) genes can cause xeroderma pigmentosum (XP), trichothiodystrophy (TTD) and sometimes combined features of XP and Cockayne syndrome (CS) (18-21). No disease mutations in other core TFIIH subunits are known, except for mutations in the *GTF2H5* (p8; also referred to as TTD-A) gene, which are known to cause TTD (22).

So far, all patient mutations found in the *ERCC3* gene are located in domains other than its two RecA like domains (HD1 and HD2), which harbor all seven conserved helicase motifs (21). A crystal structure of an archaeal homolog of XPB from *Archaeoglobus fulgidus* revealed in addition to the helicase domains the presence of a damage recognition domain (DRD) and two unique motifs named the thumb like motif (ThM) and the RED motif (4). The human XPB protein is substantially longer than its archaeal homologue and contains N- and C-terminal extensions. In order to perform its ATPase activity, XPB is suggested to bind the DNA duplex while bringing its ThM and RED motifs into close proximity (4) to each other and mutations in either motif were shown to exhibit a reduced DNA-dependent ATPase activity comparable to a mutation in the Walker A motif (K346 in human XPB) (9). However, rather than being a “true” helicase, XPB acts more likely as an ATP-dependent DNA-translocase and it was shown that its ATPase activity in transcription is utilized to track along the non-template strand in a 5’-3’ direction (23, 24). Analogous to the stimulation of XPD’s helicase activity by p44, the enzymatic activity of XPB is also regulated by its binding partner p52 (6). The importance of the interaction between XPB and p52 is demonstrated by the severe effects of an XPB mutation (F99S) found in XP/CS patients. The F99S mutation was shown to weaken the interaction between XPB and p52, resulting in a highly reduced ATPase activity of XPB (6). However, so far, it is not known how this regulation is achieved. The lack of structural information further hindered an understanding of the functional interaction between XPB and p52. Recently, cryo-EM structures of the human and yeast TFIIH complex at resolutions between 3.7 Å and 4.7 Å (25-28) were solved and shed light on the intricate interactions within TFIIH. The network between XPB and p52 is expanded by interactions to p8 and in the human TFIIH model it was shown that the TTD causing mutation L21P in p8 is located at the HD2 interface of XPB (25). How the intricate network of protein interactions regulates XPB activity is, however, poorly understood so far.

In this study, we functionally dissected the regulatory events in XPB activation utilizing proteins from the eukaryotic fungus *Chaetomium thermophilum* as a model system. We solved the crystal structure of a p52/p8 complex and used this structure in combination with the current cryo-EM models to generate point-mutations for the investigation of crucial p52/p8 interactions with XPB. We show that p52 not only acts as an activator of XPB but that also p8 in combination with p52 directly stimulates XPB’s ATPase activity further. Intriguingly, our data demonstrate that p52/p8 not only function as activators but at the same time repress the maximum activity that can be achieved when XPB is stimulated by DNA. This speed limiting can be observed in the absence of other core TFIIH components but also in the context of complete core TFIIH. Finally, we show that XPB’s ATPase activity does not directly lead to the translocase function of XPB. We show that XPB requires the presence of all core TFIIH components to act as a translocase and that the addition of XPA increases the processivity of the TFIIH translocase but not its ATPase activity. Our results suggest that XPB’s activity is tightly regulated at different levels in order to fine tune its activity for the different processes in NER and transcription.

## MATERIAL AND METHODS

### Expression and purification

The genes encoding ctXPB, ctXPD, ctp62, ctp52, ctp44, ctp34, ctp8, and ctXPA were cloned from cDNA from *C. thermophilum*. The cDNA sequence of ctXPB was codon-optimized for expression in *E. coli* (ATG:biosynthetics). CtXPB, ctp44, ctp34, ctp8, and ctXPA were inserted into the pBADM-11 vector (EMBL) containing an N-terminal hexa-histidine tag with a TEV cleavage site. CtXPD was inserted into the pBADM-11 vector without tag. Ctp62 and ctp52 were inserted into the pETM-11 vector (EMBL) without tag. The constructs ctp52_1-321, ctp52_121-514, and ctXPB_60-345 were cloned from the full length cDNA sequences via sequence and ligation independent cloning (29). The walker A variants ctXPD_K48R and ctXPB_K392R as well as all ctp52 variants were generated via site directed mutagenesis (30). CtXPB was expressed in Rosetta™ 2 (DE3) cells (Merck Millipore), ctXPD was expressed in ArcticExpress (DE3) RIL cells (Agilent). Ctp62/ctp44, ctp52/ctp8 as well as ctp52/ctp34 were co-expressed in BL21 CodonPlus (DE3) RIL cells (Agilent). Ctp8, ctp52_1-321, ctp52_121-514 and ctXPB_60-345 were separately expressed in BL21 CodonPlus (DE3) RIL cells. CtXPA was expressed in LOBSTR BL21 (DE3) RIL cells (Kerafast). For all proteins, the cells were grown in either Lennox broth or Terrific broth (Carl Roth), for the seleno-methionine containing variants of ctp52_1-321 and ctp52_121-514 the cells were grown in M9 minimal medium supplemented with L-seleno-methionine (31). For purification of the ctXPD/ctp44/ctp62 complex, ctp44 and ctp62 were co-purified via immobilized metal affinity chromatography (IMAC) using Ni IDA beads (Machery-Nagel) and size exclusion chromatography (SEC). For SEC a HiLoad 16/600 Superdex 200 prep grade column (GE Healthcare) was used with 20 mM Hepes pH 7.5, 250 mM NaCl, and 1 mM TCEP. The elution fractions containing ctp44/ctp62 were pooled and directly added to the cleared lysate of ctXPD. This mixture was subjected to another IMAC, followed by SEC and anion exchange chromatography (AEC). For SEC, the same column and buffer as above was used. For AEC a MonoQ 5/50 GL column (GE Healthcare) was used, with buffers containing 20 mM Hepes pH 7.5, 50/1000 mM NaCl, and 1 mM TCEP. CtXPB, ctXPB_60-345, ctp52/ctp34, ctp52/ctp8, ctp52_1-321, ctp52_121-514, and ctp8 were purified via IMAC (Ni TED or Ni IDA, Machery-Nagel) and SEC using a HiLoad 16/600 Superdex 200 prep grade column. For ctXPB and ctp52/ctp34, the SEC buffer contained 20 mM Hepes pH 7.5, 250 mM NaCl, and 1 mM TCEP. For ctXPB_60-345, ctp52_1-321 and ctp52_121-514, the SEC buffer contained 20 mM Tris-HCl pH 7.5, and 250 mM NaCl. For ctp52/ctp8 and ctp8 the SEC buffer contained 20 mM Hepes pH 8, and 375 mM NaCl. All variants were expressed and purified as described for the wild type proteins. In case of ctp8, two versions were used. For the co-crystallization with ctp52_121-514, ctp8 containing the N-terminal hexa-histidine tag and TEV cleavage site was used. For the other studies, the tag was removed by incubation with TEV protease after IMAC and the protein solution was subjected to another IMAC to remove the TEV protease and uncut protein followed by SEC. CtXPA was purified via IMAC and AEC using a MonoQ 5/50 GL column with buffers containing 20 mM Tris-HCl pH 8, 50/1000 mM NaCl, and 2 mM TCEP. All proteins were concentrated (50-1000 µM), flash frozen in liquid nitrogen, and stored at -80 °C. To obtain Strep-tagged XPB, ctXPB was cloned into the pFastBac vector with a C-terminal Twin-Strep-Tag and a 10x His-Tag and expressed in Hi5 cells. Protein purification was performed using a 5 ml StrepTrap HP column (GE Healthcare) followed by SEC using a HiLoad 16/600 Superdex 200 prep grade column (GE Healthcare) in a buffer containing 200 mM NaCl, 20 mM Hepes pH 8 and 1 mM TCEP. When the protein was bound to the affinity column, a high salt wash with 1 µM NaCl was performed prior to elution.

### Crystallization

Crystallization was performed via the vapour diffusion method at 20 °C. For p52_1-321, a seleno-methionine derivative was crystallized at concentrations of 4-7 mg/ml. The main reservoir consisted of 100 mM calcium acetate, 12.5 % (w/v) PEG 8000, and 100 mM HEPES pH 7.5. For p52_121-EdL, a seleno-methionine derivative was crystallized at concentrations of 5-6 mg/ml. The main reservoir consisted of 7.5 % (w/v) PEG 4000 and 100 mM HEPES pH 7.0. For the p52_121-EdL/p8 complex, p52_121-EdL at concentrations of 5-6 mg/ml was incubated for 1 h at 4 °C with a twofold excess of p8 prior to crystallization. The main reservoir consisted of 150 mM ammonium chloride, 15 % (w/v) PEG 4000, and 100 mM Tris-HCl pH 8.5.

### Structure solution and refinement

X-ray diffraction data was processed with XDS (32). Diffraction data for the p52_121-EdL/p8 complex was anisotropy-corrected using the STARANISO Server (Global Phasing Limited, http://staraniso.globalphasing.org/cgi-bin/staraniso.cgi). Initial phases were obtained using SHARP (33), utilizing the anomalous signal of the dataset from the seleno-methionine derivate of p52_121-EdL. The resulting electron density map was used to build an initial model of p52_121-EdL. This model was used to solve the phase problem for the p52_1-321 and p52_121-EdL/p8 datasets via molecular replacement with Phaser (34). The models for p52_1-321 and p52_121-EdL/p8 were manually completed and corrected with Coot (35). For the latter, the available structure of a Tfb5/Tfb2 minimal complex (PDB code 3DOM), was used as additional template for manual model building. The p52_1-321 structure was refined with REFMAC5 (36) using automated twin refinement. The p52_121-EdL/p8 structure was refined with BUSTER (Global Phasing Limited, http://www.globalphasing.com/buster/wiki/index.cgi?BusterCite).

### ATPase Assay

CtXPB’s ATPase activity was quantified utilizing an *in vitro* ATPase assay in which ATP consumption is coupled to the oxidation of NADH via pyruvate kinase and lactate dehydrogenase activities. Activities were measured at 30 °C in 100 µl solution, containing 1.3 U pyruvate kinase, 1.9 U lactate dehydrogenase (PK/LDH, Sigma), 1.6 mM phosphoenolpyruvate, 0.24 mM NADH, 10 mM KCl, 2.5 mM MgCl_2_, 1 mM TCEP, and 20 mM Tris-HCl pH 8.

The assay was carried out under saturating concentrations of ATP using ctXPB wild type and ctXPB K392R at a concentration of 250 nM. ctp52, ctp52 variants, ctp8, and ctp34 were added in a 2 fold molar excess to ctXPB. When core ctTFIIH was assembled, all subunits were present at an equimolar ratio with a final concentration of 250 nM. CtXPA was added in a 4:1 stoichiometric ratio to ctXPB or ctTFIIH, respectively. If not stated otherwise, dsDNA (fw: 5’-AGCTACCATGCCTGCACGAATTAAGCAATTCGTAATCATGGTCATAGC-3’; rv: 5’-GCTATGACCATGATTACGAATTGCTTAATTCGTGCAGGCATGGTAGCT-3’) was used with a final concentration of 250 nM for ctXPB and with a final concentration of 1 µM for ctTFIIH in order to achieve full DNA activation. The sample was preincubated until a stable base line was achieved. Enzyme catalysis was initiated by the addition of 2.5 mM ATP. The activity profiles were measured at 340 nm using a Clariostar Plate reader (BMG Labtech) and 384 well and 384 well F-bottom μClear™ microplates (Greiner Bio-One). Initial velocities were recorded, and ATP consumption was determined using the molar extinction coefficient of NADH. Curves were fitted with Graphpad Prism. All measurements were carried out in triplicates.

### Translocase Assay

Core ctTFIIH dsDNA-translocase activity was detected using a well established triplex disruption assay (24, 27). dsDNA-translocase activity is measured by displacement of a fluorescently labeled triplex forming oligonucleotide (TFO) from a triple helix DNA substrate. The assay was carried out at 30 °C in 50 µl solution containing 120 mM KCl, 8 mM MgCl_2_, 16 mM Hepes-KOH pH 8, 4 % glycerol, 0.8 mM TCEP, 2.2 mM phosphoenolpyruvate, 1.8 U pyruvate kinase (Sigma) and 150 nM triplex DNA.

The triplex DNA consists of a 52 base pair double stranded DNA with a 5’ Dabcyl label (fw: 5’-GTCTTCTTTTAAACACTATCTTCCTGCTCATTTCTTTCTTCTTTCTTTTCTT-3’; rv: 5’-Dabcyl-AAGAAAAGAAAGAAGAAAGAAATGAGCAGGAAGATAGTGTTTAAAAGAAGAC-3’) and a 5’ Cy3-tagged TFO (5’-Cy3-TTCTTTTCTTTCTTCTTTCTTT-3’). The triplex DNA was annealed as previously described (27). Correct triplex formation was checked via native PAGE. The baseline was recorded for 10 minutes prior to addition of 2 mM ATP. TFO displacement was then measured for 60 minutes at 520-540 nm excitation and 590-620 nm emission and a gain of 1900 using a Clariostar Plate reader (BMG LABTECH) and 384 well F-bottom FLUOTRAC™ high binding microplates (Greiner Bio-One). When core ctTFIIH was assembled, all subunits were present at a 1:1 stoichiometric ratio with a final concentration of 500 nM. CtXPA was added in a four-fold molar excess to ctTFIIH or ctXPB. For the minimal XPB complex, ctp52, ctp8, and ctp34 were added in a two-fold molar excess to ctXPB (500 nM). The release of Cy3-tagged TFO corresponds to the slope of increasing fluorescence. Curves were fitted with Graphpad Prism. All measurements were carried out with at least 10 replicates and with at least two different protein batches.

### Pull down assay

10 µl Strep-Tactin Sepharose beads (IBA) were mixed with 100 µl buffer containing 150 mM NaCl, 1 mM TCEP and 20 mM Hepes pH 7.5. 2 µM ctXPB with a Twin-Strep-Tag was added and incubated for 45 minutes with the beads. Then 16 µM ctp52/ctp8 wildtype or E359K were added and incubated for another 45 minutes. To exclude unspecific binding of p52/p8, 16 µM ctp52/ctp8 were added to the beads without XPB. Unbound protein was washed off and bound protein was eluted by boiling for 5 minutes at 95 °C. The samples were analyzed by SDS-PAGE. The intensities of the gel bands were quantified using ImageJ and graphs were generated via GraphPad Prism. To calculate standard deviations, three technical replicates were performed.

### Fluorescence anisotropy

DNA binding was analyzed by fluorescence anisotropy employing a splayed duplex DNA with a Cy3 label (fw: Cy3-5’-AGCTACCATGCCTGCACGAATTAAGCAATTCGTAATCATGGTCATAGC-3’ rv: 5’-GCTATGACCATGATTACGAATTGCTTGGAATCCTGACGAACTGTAG-3‘). Assays were carried out in 50 µl solution with 20 mM HEPES pH 7.5, 10 mM KCl, 2.5 mM MgCl_2_, 1 mM TCEP, and 6 nM DNA at room temperature. CtXPB was used at concentrations of 0.5 to 500 nM as indicated. Ctp52/ctp8 was added in 2:1 stoichiometric ratio to ctXPB. Fluorescence polarization was detected at an excitation wavelength of 540 nM and an emission wavelength of 590 nM with a Clariostar plate reader (BMG labtech) using 384 well F-bottom Fluotrac high binding microplates (greiner bio-one). The gain was adjusted to a well containing buffer and DNA but no protein. The measurements were carried out in triplicates. Curves were fitted with Graphpad Prism.

### CD spectroscopy

Circular dichroism (CD) spectra were recorded under constant nitrogen flush using a Jasco J-810 spectropolarimeter over 190 – 260 nm. Measurements were performed at 20 °C in 8 scans with a speed of 50 nm/min and a band width of 2 nm. The proteins were diluted to 4 µM in a buffer containing 19 mM dipotassium phosphate and 1.2 mM monopotassium phosphate at pH 8. All samples were centrifuged for 20 minutes at 25000 rpm prior to the measurement. The buffer spectrum was used as a baseline and substracted from all protein spectra.

## Results

### The ctp52/ctp8 structure

Several cryo-EM structures of TFIIH have been published recently shedding light on the molecular architecture of the TFIH components including the important p52/p8 complex (25-28). However, no full atomic model of p52, the major regulator of the XPB enzyme, alone or in complex with p8 beyond a resolution of 3.5 Å is available so far. We pursued the structural characterization of p52 from the fungal model organism *Chaetomium thermophilum* (ctp52) using x-ray crystallography. ctp52 shares 35% and 39% sequence identity with the human p52 and yeast Tfb2 sequences, respectively (Supplementary Figure 1). We solved two crystal structures of ctp52. The first structure encompasses amino acids 1-321 (ctp52_1-321), the second encompasses the C-terminal part starting at amino acid 121 (ctp52_121-514). In the latter a 28 amino acid long linker region (aa 322 to 349), that is not present in human p52, was replaced by a short linker (sequence SNGNG). Ctp52_121-514 was solved in complex with p8. The crystal structures of ctp52_1-321 and ctp52_121-514/ctp8 were solved at resolutions of 2.8 Å and 2.7 Å, respectively (Supplementary Figure 2A, B, Table 1). Due to the overlap of both structures we obtained a full structural model of ctp52 at atomic resolution (Figure 1A, Supplementary Figure 2C). The model shows that ctp52 is organized into four distinct domains, an N-terminal domain (NTD; amino acids (aa) 1-120), two middle domains (MD1; aa 121-319 and MD2; aa 350-454), and a C-terminal domain (CTD; aa 455-514), that interacts with p8 (37). The linker region that was replaced in ctp52_121-514, bridges MD1 with MD2 and could not be observed in our structure due to its flexibility. The ctp52_1-321 model comprises the NTD and MD1 domains and the ctp52_121-514 model the MD1, MD2, and CTD domains. Both crystal structures were superimposed via the MD1 domain (rmsd of 0.87 Å) thereby generating a model of full-length ctp52.

**Table 1.**
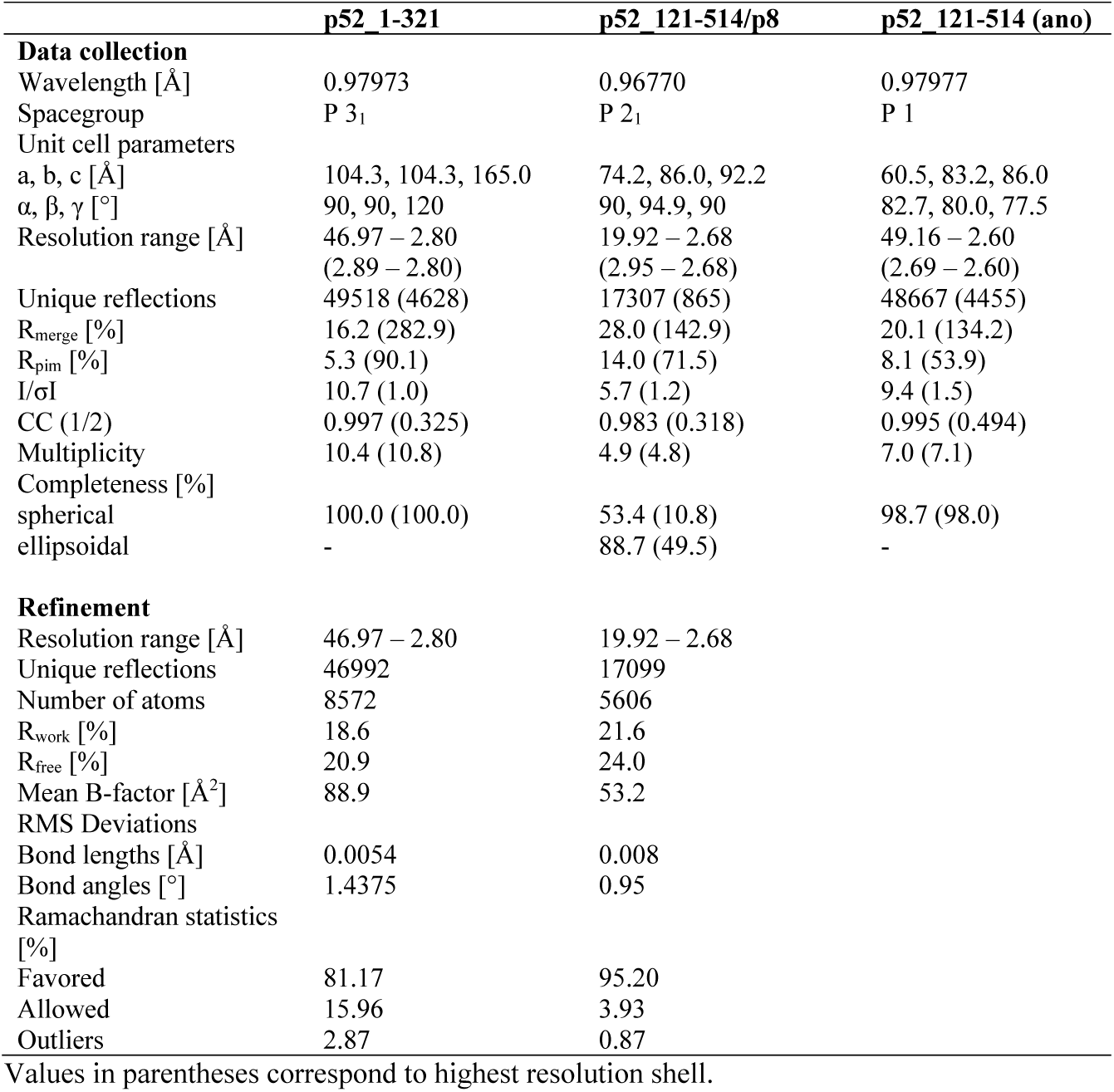
Data collection and refinement statistics

**Figure 1:**
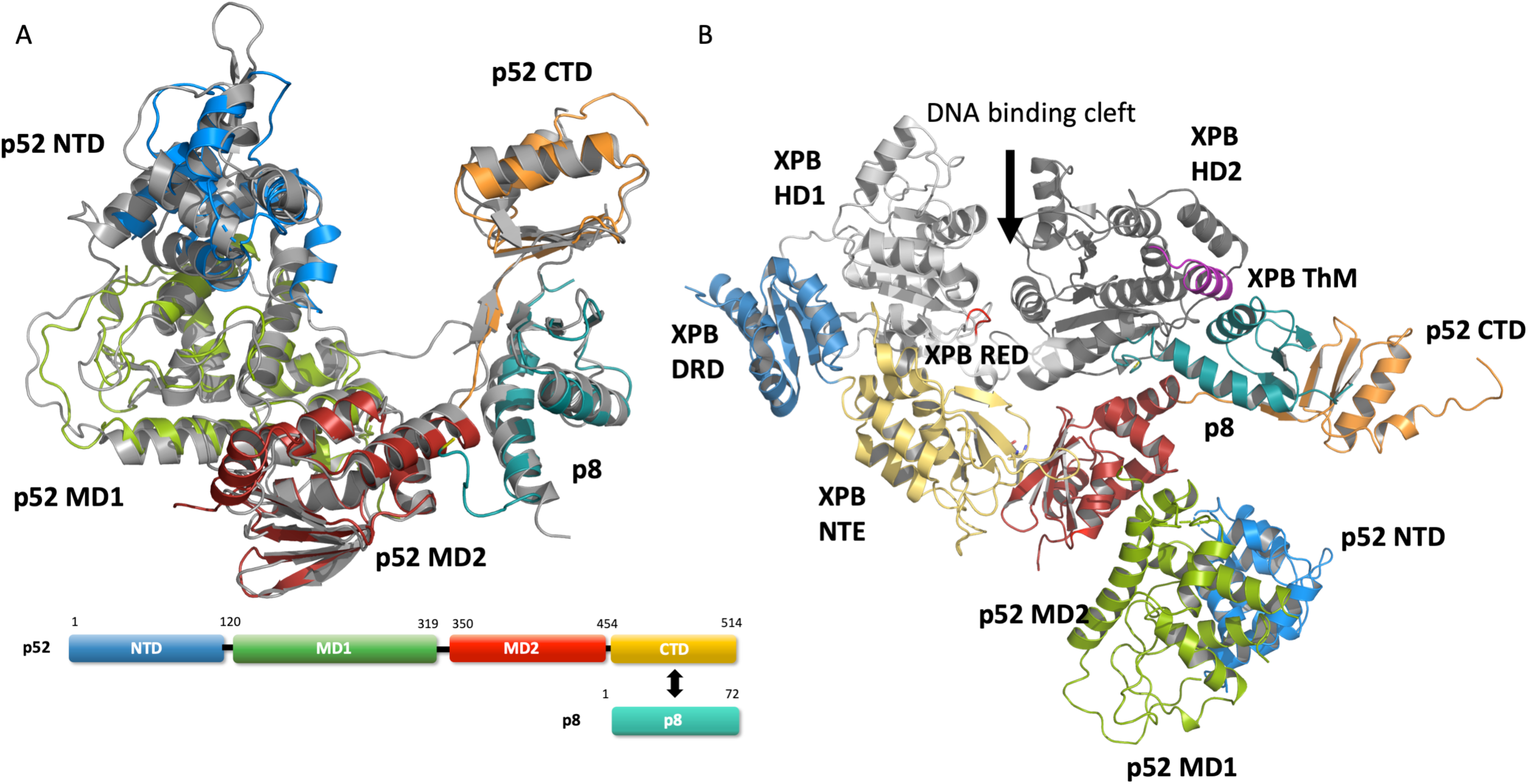
Structure and domain architecture of ctp52 and ctp8. **(A)** Structure and domain architecture of full length ctp52 and ctp8. Ctp52 is comprised of the following domains: NTD (blue), MD1 (green), MD2 (red) and CTD (orange). Ctp8 is shown in mint green. Remarkably, ctp8 and ctp52 CTD adopt the same fold. **(B)** Ctp52 and ctp8 modelled into the cryoEM structure from Greber et. al (25) in complex with XPB.

The ctp52 structure is predominantly *α*-helical and consists of 22 *α*-helices and 15 *β*-strands in total (Figure 1A, Supplementary Figure 2C). None of the newly identified domains present in ctp52 (NTD, MD1 and MD2) displays significant structural homologies to any known protein domains. The interaction between ctp52 and ctp8 is mediated by *β*-strands 12 to 15 of p52 and *β*-strands 1 to 3 of p8 and can be readily compared to the crystal structure of the minimal yeast Tfb5/Tfb2 complex (37) (PDB code: 3DOM) with an rmsd of 1.10 Å for the CTD of ctp52 and an rmsd of 0.97 Å for ctp8 (PDBsum).

A superposition of the ctp52/ctp8 structure with the cryo-EM model of human p52/p8 (25) shows that the MD2 domain was found to be oriented differently compared to our model (Supplementary Figure 2 D). While the antiparallel *β*-sheet of MD2 (*β*-strands 8-11) is pointing towards MD1 and is involved in the formation of the dimer interface in our ctp52_121-514/ctp8 structure (Supplementary Figure 2 B), it is pointing towards the N-terminal extension (NTE) of XPB in the ctp52/ctp8/XPB model (Figure 1B). This is most likely due to the different conditions and interactions in the crystallization experiments compared to the cryoEM experiment. However, the rearrangement of the individual ctp52 domains according to the model of human p52/p8 embedded in TFIIH leads to an overall rsmd of 1.6 Å indicating a high degree of structural similarity.

The main and most relevant interaction between the MD2 of p52 and the NTE of XPB seems to be mediated by an orthogonal packing of their respective *β*-sheets. Furthermore, MD2 of p52 is also interacting with the second helicase domain, HD2, of XPB in the ctp52/ctp8/XPB hybrid model (Figure 1B). This interaction involves twelve residues of the MD2 domain located mainly in helix *α*20 of p52 and the *β*-turns that connect *β*-strand 8 to 11. The other side of the interface is formed by eleven residues of XPB’s HD2. Both interfaces combined comprise an approximate surface area of 1580 Å^2^. Importantly, an extension of the interface is achieved through the presence of p8 which also interacts with the HD2 of XPB thereby providing an additional surface area of approximately 516 Å^2^ (Note that all interface surface area values were derived from 6nmi).

The hybrid structure illustrates that four entities including p8 and the MD2 of p52, together with the NTE and the damage-recognition (DRD) domains of XPB form a lunate-like ring (Figure 1B), which embraces the two RecA like domains of XPB from one side, while the DNA enters from the other side (26, 27). DNA binding, a functional Walker A motif and an intact RED (arginine-glutamate-aspartate) motif as well as the thumb-like motif (ThM) seem to be prerequisites for the activity of XPB during NER (4, 6, 9). The lunate-like ring encircles the RecA like domains from one side and thereby may stimulate XPB’s ATPase activity. Interestingly, p8 directly interacts with the ThM motif of XPB. Due to the importance of XPB’s ATPase activity in transcription and DNA repair we pursued further studies to decipher the functional influence of the individual components and interactions of this intricate network.

### The ATPase activity of ctXPB is sequentially stimulated by ctp52 and ctp8

To investigate which components are necessary for the full activation of XPB we used the proteins from *C. thermophilum* due to their superior stability. We first performed titration experiments using a co-purified ctp52/ctp34 complex. ctp34 was used to enhance the stability of ctp52 and to ensure a native ctp52 conformation in accordance with the composition in TFIIH. ctp52/ctp34 activate ctXPB in a concentration dependent manner, reaching a V_max_ of 9.3 µmol ATP*l^-1^*min^-1^ and an EC_50_ value of 178 nM (Figure 2A, open circles). When ctp8 is added to the ctXPB/ctp52/ctp34 complex the basal activity level is 7.8 µmol ATP*l^-1^*min^-1^ and reaches a maximum of 20.8 µmol ATP*l^-1^*min^-1^ as highest activity with an EC_50_ value of 310 nM (Figure 2A, open triangles) indicating that ctp8 is directly able to further enhance the ctp52-mediated activation of ctXPB. We then investigated whether a co-purified ctp52/ctp8 complex alone is able to activate ctXPB and observed a full activation with 21.1 µmol ATP*l^-1^*min^-1^ and an EC_50_ value of 159 nM (Figure 2A, open squares), showing that ctp52/ctp8 is sufficient for ctXPB activation and that ctp34 does not alter the activation cascade.

**Figure 2:**
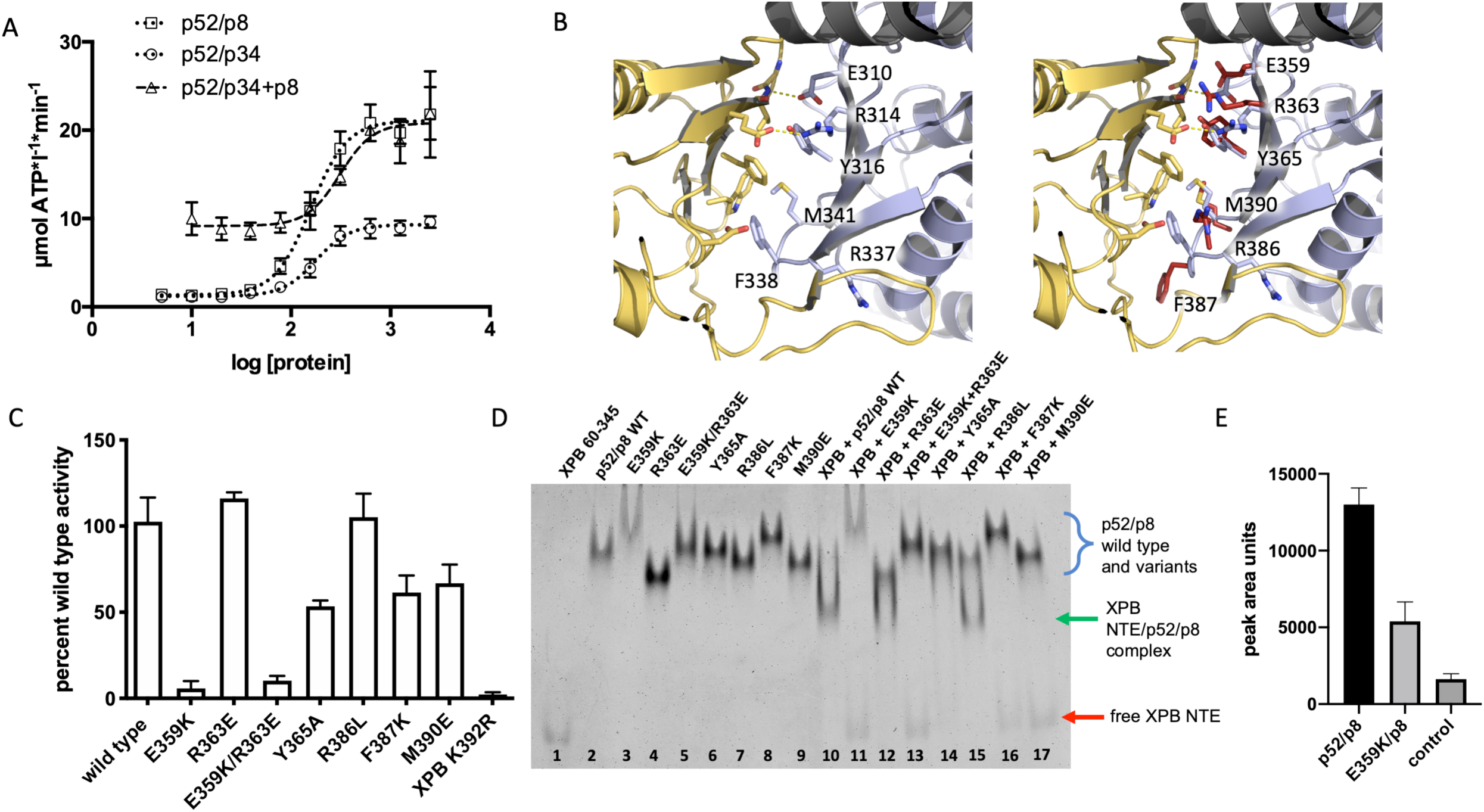
Interaction studies of ctp52 MD2 and ctXPB NTE. **(A)** ATPase activity profile of ctXPB activated by increasing amounts of different ctp52 complexes. Ctp52 complexes were titrated to a fixed amount of ctXPB. The x-axis shows the logarithmic concentration of the respective ctp52 complex. **(B)** Structural insight into the human p52 (light blue) – XPB (yellow) interface taken from 6nmi. The superpositions of ctp52 residues is shown on the right in red. The depicted ctp52 residues were mutated and analyzed with respect to their influence on ctXPB’s ATPase activity **(C)** and in terms of their interaction with the ctXPB NTE **(D). (C)** Effect of ctp52/ctp8 wild type and variants on the ATPase activity of ctXPB. Ctp52_E359K/ctp8 leads to a strong reduction of ctXPB’s ATPase activity. **(D)** Native PAGE analysis reveals loss of interaction between certain ctp52/ctp8 variants and ctXPB NTE (ctXPB 60-345). Individual proteins are shown in lanes 1-9. Ctp52 wildtype and variants in complex with ctp8 and ctXPB 60-345 are shown in lanes 10-17. **(E)** Quantification of ctp52/ctp8 and ctE359K/ctp8 pull down experiments using Streptavidin tagged ctXPB as bait. Data originate from three technical replicates. SDS PAGES were analyzed with ImageJ for quantification and mean values with SD were derived with GraphPad Prism.

Based on our ctp52 crystal structure modelled into the most recent cryo-EM map we were able to identify conserved ctp52 residues in the MD2 domain that interact with the ctXPB NTE and chose them for functional mutagenesis studies since it was also shown that some of these residues (E359, R363) led to phenotypic consequences when they are mutated (38). Therefore, we generated the following point mutations: E359K, R363E, E359K/R363E, Y365A, R386L, F387K, and M390E (Figure 2B). All variants are located in a *β*-sheet that interacts with a similar motif within XPB. We thus targeted the entire interface from the ctp52 side ranging from a hydrophobic area comprised of Y365, F387, and M390 to the charge complementary patch constituted by E359 and R363. R386 displays different side chain conformations in the cryo-EM compared to the crystal structure and was thus chosen to assess which conformation is relevant (Figure 2B). All p52 variants were subjected to CD-spectroscopy to ensure that no misfolded variants were used in the analysis (Supplementary Figure 3). Based on the observation that ctp52/ctp8 leads to a higher activation of XPB’s ATPase than ctp52 alone we investigated whether those variants were still able to activate ctXPB in the ctp52/ctp8 context (Figure 2C). The most severe impact on ctXPB activation was observed for the E359K variant that was only able to activate ctXPB to 6% of the wild type level, whereas the adjacent R363E variant had no effect and displays 116% of wild type activity. In line with the result for the single E359K variant the E359K/R363E variant displays a comparable decrease leading to remaining 10% activation. The Y365A, F387K, and M390E variants all display a medium phenotype with an activation profile of 54%, 61%, and 67%, respectively, indicating at least partial activation. Mutation of R386 to a leucine (R386L) has no effect with an activation activity of 105% compared to the wild type proteins. To further investigate whether these variants still interact with the NTE of ctXPB we performed native PAGE analysis using the ctp52 variants in complex with ctp8 and the NTE of ctXPB comprising residues 60-345 (ctXPB_NTE, Figure 2D). All ctp52/ctp8 variant complexes form a single band in the native PAGE, however, due to the different charge properties of the variants they are located at slightly different heights (Figure 2D, lanes 3-9) compared to the wild type complex (Figure 2D, lane 2). Combining ctXPB_NTE with wild type ctp52/ctp8 clearly leads to complex formation (Figure 2D, lane 10), whereas the variants that display reduced activation capabilities also show no interaction in the native PAGE experiment (Figure 2D, lanes 11, 13-14, 16-17). Since the cryo-EM structure of holo-TFIIH suggests the presence of a second XPB-ctp52/ctp8 interface located in the HD2 domain of XPB and the MD2 of p52 (Figure 1B) (25) we analyzed the E359K variant towards its interaction with full length ctXPB which indeed led to decreased but significant complex formation (Figure 2E). Importantly, this result suggests that disrupting one of the interfaces is sufficient to prevent p52/p8 mediated XPB activation, whereas complex formation is maintained.

### ctp52/ctp8 limits ctXPB DNA dependent ATPase activity

We next aimed to investigate the influence of DNA binding on the ATPase activity of ctXPB. In the sole presence of dsDNA the activity of ctXPB reached a V_max_ of 65 µmol ATP*l^-1^*min^-1^ (Figure 3A). Interestingly, this is about three times higher than the activation of ctXPB via ctp52/ctp8 (See Figure 2A for comparison). Surprisingly, in the presence of ctp52/ctp8 and dsDNA the induced ATPase activity is limited to 19 µmol ATP*l^-1^*min^-1^ which corresponds well to the activity achieved with ctp52/ctp8 in the absence of dsDNA (Figure 2A). This indicates that ctp52/ctp8 negatively controls the dsDNA-dependent activation of XPB which is illustrated further when we titrated increasing amounts of ctp52/ctp8 to an equimolar ctXPB/DNA complex. At 250 nM dsDNA ctXPB is already completely activated with an activity of 59 µmol ATP*l^-1^*min^-1^ (Figure 3A). This activity decreases in a ctp52/ctp8 concentration dependent manner until it reaches again 18.8 µmol ATP*l^-1^*min^-1^ with an IC_50_ value of 137 nM (Figure 3A), which indicates high binding affinity between ctXPB and ctp52/ctp8 and once an equimolar complex is formed downregulation is complete. To further decipher the mechanism of activity limitation we investigated whether the binding of ctp52/ctp8 influences the DNA binding capacity of ctXPB. We therefore pursued fluorescence anisotropy measurements comparing ctXPB alone and the ctXPB/ctp52/ctp8 complex (Figure 3B). The obtained K_D_ values for ctXPB of 78 nM and for the ctXPB/ctp52/ctp8 complex of 51 nM are similar indicating that there is no major effect on DNA binding caused by the presence of ctp52/ctp8. Since DNA binding did not seem to be affected, we evaluated whether ATP binding is altered by the presence of ctp52/ctp8. Michaelis-Menten experiments to assess the apparent K_M_ for ATP (Figure 3C) led to K_M_ values of 62 µM (ctXPB/DNA), 154 µM (ctXPB/ctp52/ctp8 complex) and 45 µM (ctXPB/ctp52/ctp8 complex with excess dsDNA). These values indicate a positive effect on the K_M_ for ATP in the presence of DNA. However, the corresponding V_max_ value of ctXPB with excess dsDNA (74 µmol ATP*l^-1^*min^-1^) is three times higher than V_max_ of ctXPB/ctp52/ctp8 plus DNA (23 µmol ATP*l^-1^*min^-1^). Again, the latter value fits well with the V_max_ of ctXPB/ctp52/ctp8 without DNA (24 µmol ATP*l^-1^*min^-1^) indicating that ctp52/ctp8 predominantly affects the V_max_ of the ctXPB/DNA complex. To further substantiate these observations, we investigated whether the ctp52/ctp8 complex containing the ctp52 E359K variant which abrogates the interaction with the ctXPB_NTE and does not activate XPBs ATPase activity but still permits the interaction via the HD2 of ctXPB can repress the ctXPB DNA activation (Figure 3D, Figure 2E). Our data clearly show that the V_max_ of the ctXPB/ctp52_E359K/ctp8 complex is not repressed and reaches ∼80% of wild type activity; ctp8 by itself also does not exert any effect. To assess the influence ctp8 might have on the DNA activation mechanism we titrated DNA onto ctXPB/ctp52/ctp34 and ctXPB/ctp52/ctp34/ctp8 complexes and compared them to the DNA activation of ctXPB alone (Figure 4A). In the absence of ctp8 the complex is activated two-fold by DNA from 8 to 17 µmol ATP*l^-1^*min^-1^ whereas the complex with ctp8 displays only minor activation from 18 to 22 µmol ATP*l^-1^*min^-1^. The latter two values are very similar to the maximum level of activation reached by ctp52/ctp8 in the presence or absence of DNA (23 µmol ATP*l^-1^*min^-1^). It seems that ctp8 in this case partially compensates for the presence of DNA since the maximum activity value of the ctXPB/ctp52/ctp34 complex in excess of DNA (17 µmol ATP*l^-1^*min^-1^) is similar to the value of the ctXPB/ctp52/ctp34/ctp8 complex (22 µmol ATP*l^-1^*min^-1^). It is important to note, however, that none of these complexes reached the rate of DNA induced activity in the sole presence of ctXPB (Figure 4A).

**Figure 3:**
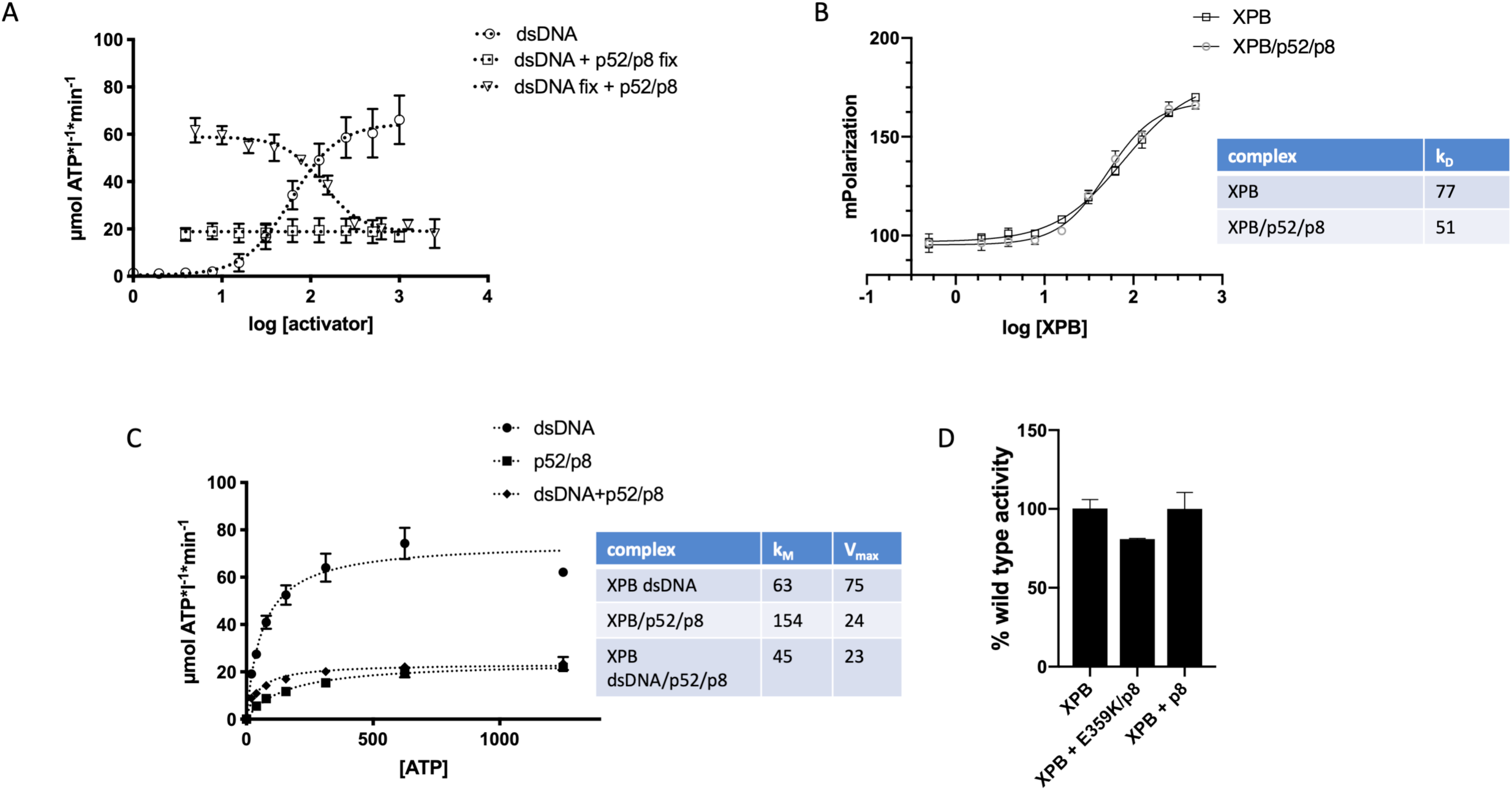
Influence of DNA and nucleotide binding on ctXPB’s ATPase activity. **(A)** Influence of DNA and ctp52/ctp8 on the ATPase activity of ctXPB. Increasing amounts of DNA were titrated onto fixed concentrations of ctXPB (circles) and ctXPB with ctp52/ctp8 (squares). Increasing amounts of ctp52/ctp8 were titrated onto fixed concentrations of ctXPB plus DNA (triangles). The x-axis shows the logarithmic concentration of the respective complex that was titrated. **(B)** Fluorescence anisotropy measurements to investigate the influence of ctp52/ctp8 on ctXPB’s dsDNA binding ability. Increasing amounts of ctXPB (square) and ctXPB with ctp52/ctp8 (circle) were titrated onto fixed amounts of fluorescently labeled DNA. Ctp52/ctp8 was used in 2:1 stoichiometric ratio relative to ctXPB. The indicated protein concentrations refer to ctXPB. **(C)** ATPase activity profile with Michaelis Menten kinetics. Increasing amounts of substrate (ATP) were titrated onto different complexes of ctXPB with its activators DNA (rhomb), ctp52/ctp8 (square) and DNA plus ctp52/ctp8 (circle). Table: V_max_ and K_M_ values for different ctXPB activation complexes. **(D)** ATPase activity of ctXPB in the presence of DNA, ctXPB in the presence of ctp52-E359K/ctp8 and DNA or just ctXPB with ctp8 and DNA.

**Figure 4:**
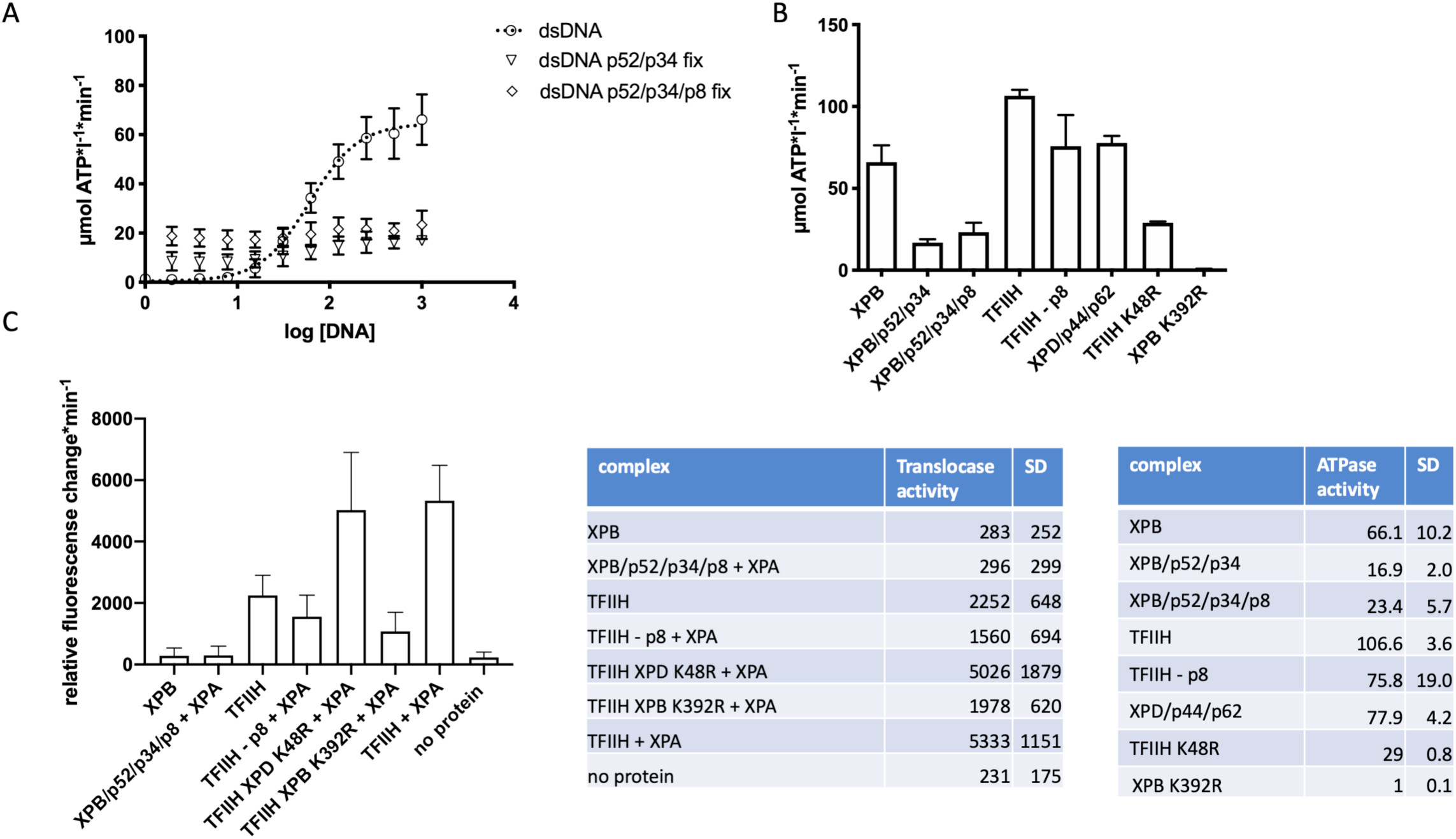
ATPase and translocase activities of ctXPB in core ctTFIIH. **(A)** ATPase activity profile of ctXPB activated by increasing amounts of DNA without ctp52 (circle) and with fixed concentrations of ctp52/ctp34 (triangle) and ctp52/ctp34/ctp8 (rhomb). **(B)** ATPase activity of core ctTFIIH and subunit complexes in the presence of 1 µM DNA. K48R indicates the catalytically dead ctXPD variant. **(C)** dsDNA translocation activity of core ctTFIIH and different subunit complexes in the presence and absence of ctXPA. In all instances where TFIIH is mentioned in the panels this refers to core TFIIH.

We further investigated the activity of ctXPB in the broader context of core ctTFIIH by adding a ctXPD/ctp44/ctp62 complex to ctXPB/ctp52/ctp34/ctp8. In core ctTFIIH, the overall dsDNA (1µM) induced ATPase activity is higher than observed for ctXPB alone with 107 µmol ATP*l^-1^*min^-1^ compared to 66 µmol ATP*l^-1^*min^-1^, respectively (Figure 4B). We showed that the ctXPB/ctp52t/ctp34 and ctXPB/ctp52/ctp34/ctp8 complexes in the presence of DNA display low activities of 17 µmol ATP*l^-1^*min^-1^ and 23 µmol ATP*l^-1^*min^-1^, respectively (see above), whereas the ctXPD/ctp44/ctp62 complex displays an ATPase activity of 78 µmol ATP*l^-1^*min^-1^. The activities of both helicase complexes thus add up to 101 µmol ATP*l^-1^*min^-1^ which is almost identical to our observed core ctTFIIH ATPase activity indicating that no further activation of the ctXPB ATPase in the core ctTFIIH environment is achieved by the presence of the additional subunits. This is in line with the core ctTFIIH XPD K48R variant (XPD Walker A mutant) and an observed activity of 29 µmol ATP*l^-1^*min^-1^. Core ctTFIIH lacking ctp8 displays a reduction in activity of around 30 µmol ATP*l^-1^*min^-1^ (76 µmol ATP*l^-1^*min^-1^). This reduction in activity is higher than expected based on our results obtained with the ctXPB/ctp52/ctp34 and the ctXPB/ctp52/ctp34/ctp8 complexes, which emphasizes the importance of ctp8 within core ctTFIIH (39) and indicates that XPB’s ATPase activity is mainly regulated by p52/p8 but not by other subunits within TFIIH.

### ctXPB translocase activity is dependent on core ctTFIIH

After assessing how ctXPB’s ATPase function is regulated, we investigated how this regulation affects the translocase function of ctXPB. XPB is the essential enzymatic component of the TFIIH 5’-3’ translocase complex (24), therefore its ATPase activity should directly influence its ability to translocate on dsDNA. We used a well established triplex disruption assay to monitor ctXPB translocase activity (24, 27). Reconstituted core ctTFIIH exhibits a robust translocase activity (2250 u rel. FC*min^-1^) that is further activated by the addition of a 4-fold molecular excess of ctXPA (5333 u rel. FC*min^-1^, Figure 4C). Reconstituting the core ctTFIIH/ctXPA complex with the ctXPD walker A mutant has no effect on the translocase activity (5026u rel. FC*min^-1^) whereas reconstituting the core ctTFIIH/ctXPA complex with the ctXPB walker A mutant (ctXPB K392R) shows a strong decrease in activity (Figure 4C). These data clearly show that ctXPB is the major motor for translocase activity within core ctTFIIH. Interestingly, removal of ctp8 from the core ctTFIIH/ctXPA complex also decreases the activity significantly to 1560u rel. FC*min^-1^. To our surprise ctXPB by itself does not exhibit any measurable translocase activity within our experimental conditions (283u rel. FC*min^-1^, which is comparable to the negative control of 231u rel. FC*min^-1^). The formation of “half” of core TFIIH through the assembly of ctXPB with ctp52/ctp34/ctp8 and a 4-fold excess of ctXPA also yields no measurable translocase activity (296u rel. FC*min^-1^). To investigate, if the increase in translocase activity by ctXPA is due to the stimulation of XPB’s ATPase activity we performed ATPase assays. However, ctXPA does not seem to influence ctXPBs ATPase activity in any of the tested conditions (Suppl. Figure 4), thus excluding the possibility that the translocase effect mediated by XPA is due to further ATPase modulation. Our data therefore suggest that the XPB translocase function is only operating in the context of core TFIIH and the enzymatic activity is solely provided by XPB.

## Discussion

Recent structural data derived from cryo-EM studies have greatly improved our understanding towards the architecture of core TFIIH and its individual components (25-27). However, little is known so far about the intricate activity network within core TFIIH and how the individual subunits define and regulate the activity of the entire complex. In this work we focused on the XPB helicase and investigated the regulation of XPB (i) by its direct interaction partners p52 and p8; and (ii) in the context of the entire core TFIIH, to obtain insights how the ATPase function is regulated, since it is essential for transcription and DNA repair.

Our crystal structures of the ctp52/ctp8 complexes show that p52 is a flexible protein that requires its binding partners XPB and p8 to assume a complete functional state. The high structural homology of the individual p52 domains but the different orientations of the domains relative to each other within our crystal structures and the cryo-EM TFIIH structures clearly provides support for this hypothesis. The modular composition of p52, however, may also be necessary to provide the flexibility to assume different conformational states to fulfill TFIIH’s functions in transcription and NER (25-27, 40). Importantly, the high structural homology between the *C. thermophilum* and the human proteins indicates a high functional homology of the two organisms that has been described and utilized previously (11, 41). In this study, we extended the utilisation of this system towards the analysis of XPB regulation and the functional assembly of core TFIIH. We initiated our analysis towards the interaction of ctXPB and ctp52 and the effect that ctp52 exerts on ctXPB. We observed that ctp52 is sufficient to enable XPBs ATPase function (9.3 µmol ATP*l^-1^*min^-1^), which is well in line with previous work (6). To our surprise, this effect could be directly boosted by the addition of ctp8 (20.8 µmol ATP*l^-1^*min^-1^) without the presence of other core TFIIH proteins indicating a cooperative activation mechanism depending on p52/p8 only (Figure 2A). One of the crucial players towards mediating the effect of p52 activation is the NTE of XPB. We analyzed the interface formed by the NTE of XPB and MD2 of p52 through functional mutagenesis studies and showed that individual residues within this interface are critical for the p52 dependent ATPase activation. Based on the intricate network of interactions between XPB, p52, and p8 as observed in the cryo-EM structures, a sequential activation mechanism of XPB could be envisioned (Figure 5). In its apo form XPB is not able to hydrolyse ATP due to the high flexibility of the RecA like Helicase domains (HD1 and HD2, Figure 5B), as observed in the crystal structures of the isolated XPB protein in which the helicase domains assume an orientation relative to each other which is not productive for ATP hydrolysis (4). Stabilisation and correct positioning is achieved by the combined action of p52 and p8. P52 interacts with the NTE and HD2 of XPB thereby positioning HD1 and HD2 in an ATP hydrolysis competent state (25). Stability is further enhanced by p8, which binds to an additional region in HD2 thereby resulting in further activation of the ATPase activity of XPB (Figure 5B). However, when we analysed the interaction of XPB in the presence of dsDNA as an activator we observed that the activity is about three times higher than with p52/p8 indicating that higher ATPase rates of the XPB scaffold can be achieved. To our surprise, the combination of dsDNA and p52/p8 did not lead to cooperative activation. On the contrary, the presence of p52/p8 limits the activation of XPB to the state that was achieved just by the addition of p52/p8 (Figure 3A) which could be attributed to limiting the V_max_ value of the ATPase function and not to DNA binding or changes of the apparent K_M_ for ATP (Figure 3B,C). Thus p52/p8 acts not only as a clutch (26) but also as a ‘speed limiter’ ensuring that XPB does not proceed too fast either in the presence or absence of DNA (Figure 5B). Remarkably, this speed limitation is also not released in the presence of the other core TFIIH subunits (Figure 4B) indicating that the regulatory network depicted in Figure 5A is sufficient for the incorporation and regulation of XPBs ATPase within the TFIIH scaffold. This mechanistic control requires the interactions of the NTE and HD2 domains of XPB with the MD2 of p52. Disrupting the contact points between the NTE of XPB and MD2 of p52 leads to the abrogation of the control of XPB by p52/p8 and restores the activation of XPB by DNA (Figure 2C, 3D). It thus seems that the presence of p52/p8 on the one hand is required for XPB to assume an ATPase competent conformation but on the other hand restricts that conformation so that higher ATPase rates cannot be achieved and thereby also assumes the role as a speed limiter.

**Figure 5:**
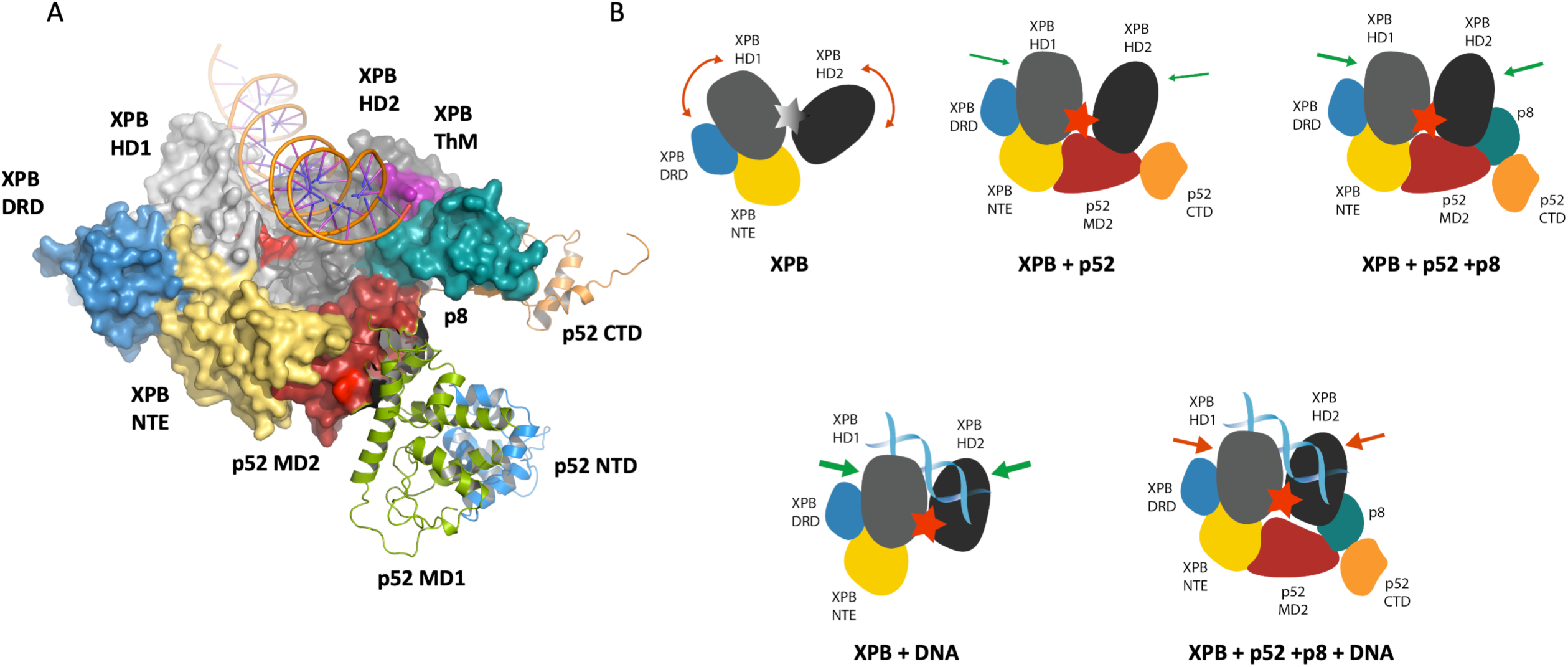
Structure and conformational changes of XPB upon activation. **(A)** Structural model of the lunate-like ring that encircles the two XPB helicase domains. Human XPB (hsXPB) comprises the following domains: NTE (yellow), DRD domain (blue), HD1 (light grey) and HD2 (dark grey) with the ThM (purple) and RED (red) motifs, respectively. The ctp52 domains MD2 (red), MD1 (green), NTD (blue) and CTD (orange) and p8 (mint green) are derived from our crystal structure. The hsXPB structure and the DNA are derived from Schilbach, *et al*. (26). The DNA duplex is modeled in orange. The lunate-like ring reaches from XPB DRD and NTE to p52 MD2 and p8 bringing the two helicase domains of XPB into close proximity and thus enabling ATP hydrolysis. P8 further expands the lunate-like ring. The ctp52 subunits that do not participate towards the formation of the lunate-like ring are modeled as cartoon. **(B)** Model of the differential XPB activation by DNA, p52/p8, p52 or p52/p8/DNA. If no activator is present, the XPB helicase domains HD1 and HD2 are too flexible and ATP hydrolysis cannot be performed. P52 binding fixes the helicase domains and thus brings them into close proximity to each other thereby enabling ATP hydrolysis. Binding of p8 further extends the lunate-like ring and further restrains the flexibility of the two helicase domains resulting in a higher activity. When DNA binds to XPB, the helicase domains are also restrained and brought into close proximity thereby enabling ATP hydrolysis leading to the highest ATPase activity. Simultaneous p52 or p52/p8 and DNA binding also results in an activation of XPB, but the activation level is limited due to the reduced flexibility of XPB.

The ATPase function of XPB is a prerogative for its subsequent actions in transcription and DNA repair (reviewed in (42)) and highly associated with its translocase activity which is involved in promoter opening and might also be important for NER (24, 27, 43). We investigated the different ATPase competent XPB complexes in a translocase assay and to our surprise the ATPase function of XPB alone is not sufficient for measurable translocase activity (Figure 4C). XPB by itself and a complex comprised of XPB/p52/p34/p8/XPA did not display any translocase activity. Only in a fully assembled core TFIIH we could observe translocase activity which is exclusively dependent on the ATPase activity of XPB, as shown by the comparison of core TFIIH assembled with the Walker A motif variants of XPD and XPB.. XPA was able to boost the translocase activity about 2.4-fold which is in line with the data from Kokic *et al* (27). XPA interacts with dsDNA above the XPB subunit of core TFIIH like a clamp, thereby restraining XPB (27). This conformation seems to stimulate the translocase activity of core TFIIH. Intriguingly, this stimulation is not caused by a modulation of the ATPase activity of XPB as shown in Supplementary Figure 4 and we thus suggest that XPA increases the processivity of the complex. Interestingly, despite its size, p8 plays a highly important role with respect to the translocase activity within core TFIIH. Removal of p8 from core TFIIH causes a significant decrease in XPA stimulated translocase activity that is even lower than the activity of core TFIIH without XPA (Figure 4C).

The impact of p8 on XPBs translocase activity is twofold, it’s loss leads to a reduction in ATPase activity but it also suggests that XPA recruitment could be impaired (44) which subsequently further diminishes the translocase activity due to the lack of XPA stimulation. Combined, these observations emphasize the role of p8 as a vital factor for NER and transcription as reflected by its involvement in the disease TTD (39, 45).

Our data show that XPB is a highly regulated protein and this regulation is embedded in several layers. The activation of XPB by DNA may only be important for other, less explored functions of XPB related to its localisation at the centrosome during mitosis or mRNA export (46, 47) and thus requires different regulation or possibly even higher activity than in the context of TFIIH. When XPB fulfils its essential functions in transcription and NER the main regulators are p52 and p8 that are able to activate XPB in a cooperative manner and at the same time act as a speed limiter, seemingly keeping the ATPase activity always at a constant level.

XPB’s ATPase activity is at the core of NER and transcription initiation (48), however, the requirements for XPB in both processes differ. In transcription the absence of XPB is less detrimental than its presence in an inactive state (49). Since XPB exhibits ATPase activity instantaneously when combined with p52/p8 no additional trigger is required in the TFIIH environment rendering XPB the only functional ATPase in a non-DNA bound TFIIH, whereas XPD activity is strictly DNA-dependent (11). This DNA independent ATPase activity could be instrumental to rearrangement/recruitment processes in the initial steps of transcription and then result in a translocation dependent removal of a block imposed by XPB itself (49). This hypothesis is also supported by the low translocase activity of core TFIIH in the absence of XPA. In NER XPB’s ATPase is essential and a DNA independent ATPase activity could aid towards the initial steps of TFIIH recruitment by XPC (9) and subsequent engagement with DNA. After these initial steps the arrival of XPA would stimulate the translocase activity to enhance the unwinding of the repair bubble necessary for XPD to engage with the DNA, yielding a repair competent complex that would also be in line with XPA’s role in lesion verification (50).

In this work, we have established novel concepts of how the XPB ATPase is regulated by the intricate interaction network within TFIIH. We show in a stepwise approach how p52 and p8 are able to activate XPB in an additive way, importantly, this is independent from the activation of XPB by DNA. Despite being seen as an activator the p52/p8 complex also limits the V_max_ of the XPB ATPase activation by DNA and thus assumes the function of a speed limiter clearly dominating regulation within TFIIH. Lastly, we show that despite its full ATPase capacity in isolation XPB only acts as a translocase in the context of core TFIIH. Combined, our work redefines the regulatory network of the XPB ATPase and translocase shedding light into the inner regulatory mechanisms of core TFIIH in transcription and repair.

## Data Availability

Atomic coordinates and structure factors have been deposited in the Protein Data Bank under the accession codes 6trs (ctp52_121-514/p8) and 6tru (ctp52_1-321).

## Supplementary Data statement

Supplementary Data are available at NAR Online.

## Acknowledgments

We would like to thank the staff from the beamline ID30A-3 at the European Synchrotron Radiation Facility and the staff at beamline 14.1 at BESSY for excellent support.

