## Supplementary Data for "How to limit the speed of a motor: The intricate regulation of the XPB ATPase and Translocase in TFIIH"

**Supplementary Figure Legends:**

**Supplementary Figure 1: Multiple sequence alignment of p52 MD2.** Protein sequences of p52 MD2 from *Homo sapiens*, *Chaetomium thermophilum*, *Saccharomyces cerevisiae* and *Drosophila melanogaster* were aligned using T-coffee (51) and ESPript 3.0 (52). The p52 variants generated in this study are highlighted in blue and are highly conserved. Amino acid numbers are shown for the human protein.

**Supplementary Figure 2: Crystal structure of full length ctp52 in complex with ctp8.** **(A)** Crystal structure of the ctp52_1-321 construct that comprises the NTD and MD1 of ctp52. There are two molecules in the asymmetric unit, the second molecule is shown in grey. **(B)** Crystal structure of the ctp52_121-514 construct that comprises MD1, MD2 and CTD of ctp52, as well as ctp8. The TEV helix is involved in dimerization. There are two molecules in the asymmetric unit, the second molecule is shown in grey. **(C)** Superposition of the two crystal structures from (A) and (B) via MD1 generates the full length ctp52 model. **(D)** Superposition of our ctp52 structure with the human p52 and p8 structure in the context of TFIIH (25). In the TFIIH cryo-EM structure MD2 is twisted and the orientation of p52 CTD and p8 is different, compared to our structure.

**Supplementary Figure 3: CD spectroscopy of ctp52 wild type and variants.** Analysis of ctp52 wild type and variants in complex with ctp8 via CD spectroscopy to assess whether they all assume the same fold.

**Supplementary Figure 4: ATPase activity of different ctTFIIH subunit complexes in the presence and absence of ctXPA and DNA.** Different ctTFIIH complexes were analyzed in the presence and absence of DNA as well as with and without ctXPA. Neither the addition of DNA, nor the addition of ctXPA has any influence on the ATPase activity of ctTFIIH or the subunit complexes.

### Supplementary Figures:

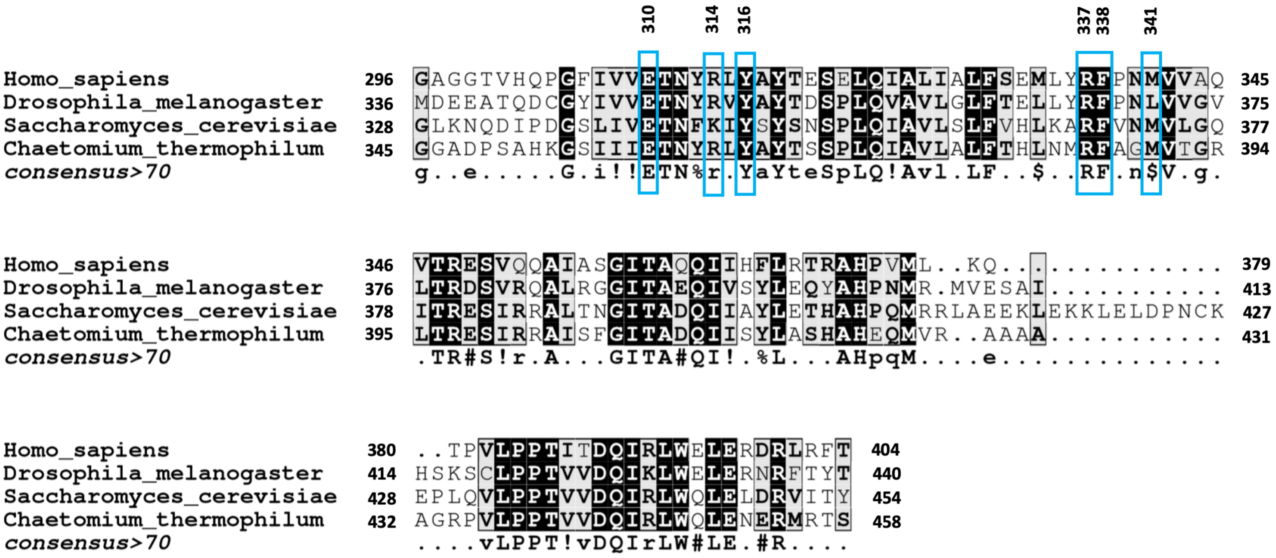


Supplementary Figure 1

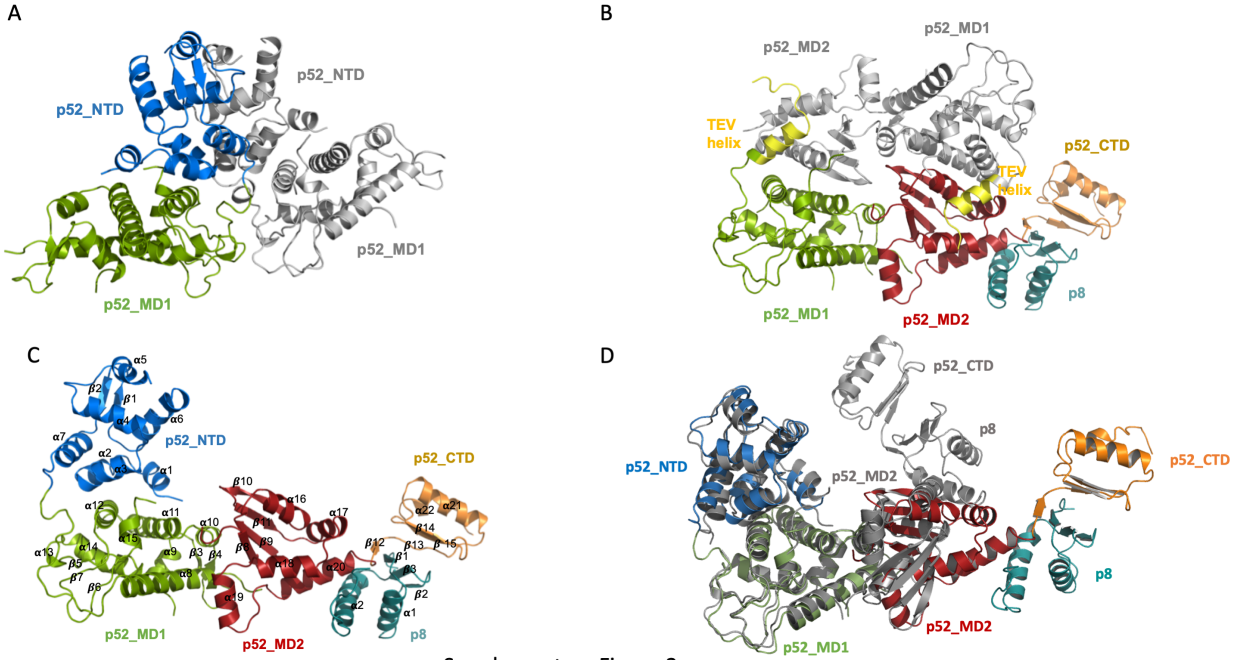


Supplementary Figure 2

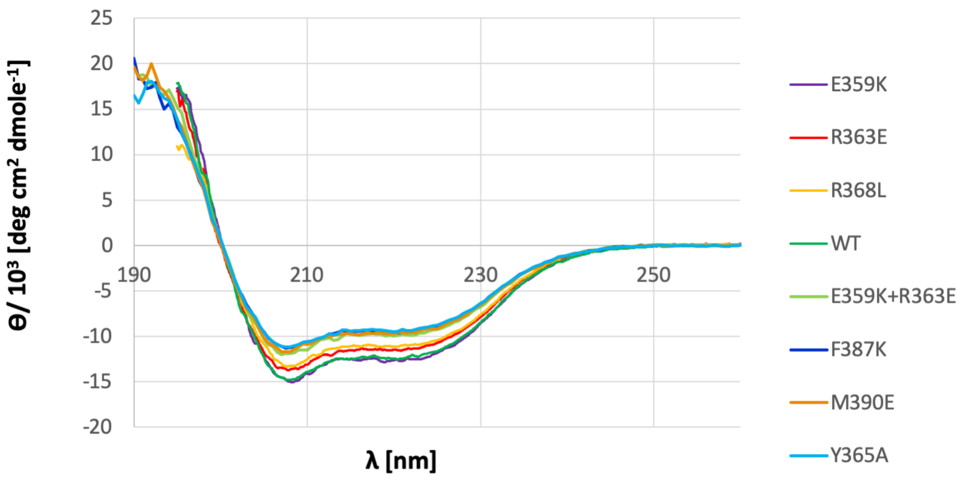


Supplementary Figure 3

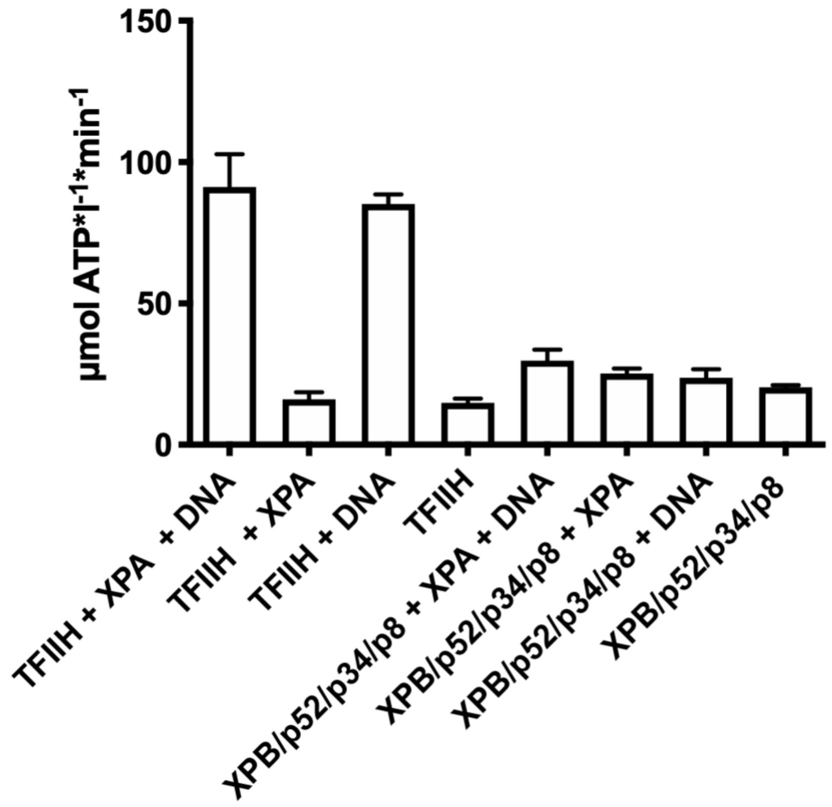


Supplementary Figure 4
